# Impact of Paraben on Uterine Collagen: An Integrated and Targeted Correlative Approach Using Second Harmonic Generation Microscopy, Nanoindentation, and Atomic Force Microscopy

**DOI:** 10.1101/2024.11.06.622338

**Authors:** Mahmuda R. Arshee, Ritwik Shukla, Jie Li, Indrani C. Bagchi, Ayelet Ziv-Gal, A. J. Wagoner Johnson

## Abstract

This study investigates the structural and mechanical changes in uterine collagen following exposure to propylparaben (PP), using a combined methodology of Second Harmonic Generation (SHG) microscopy, Nanoindentation (NI), and Atomic Force Microscopy (AFM). SHG analysis identified significant disorganization in collagen fibril orientation in the circumferential layer and heterogeneous distribution of regions with elevated forward to backward ratios (F/B) across all uterine layers due to PP exposure. High F/B can indicate multiple potential fibril-level changes like thickened fibrils, higher crosslinking, fibril disorganization - changes not fully decipherable by SHG alone. Recognizing this limitation, the study employs NI and AFM to provide complementary mechanical and nanoscale insights. NI revealed increased indentation modulus in the exposed uteri, suggesting increased stiffness. Co-registration of the indentation response with SHG parameters uncovered that elevated F/B regions show enhanced mechanical stiffness, suggesting a fibrotic transformation following PP exposure. AFM was specifically performed on regions identified by SHG as having low or high F/B, providing the necessary nanoscale resolution to elucidate the structural changes in fibrils that are likely responsible for the observed alterations. This approach confirmed the presence of disordered and entangled collagen fibrils in the circumferential layer in all regions and an increase in fibril diameter in the high F/B regions in the exposed uteri. Together, these findings demonstrate significant alterations in collagen architecture due to PP exposure, revealing disruptions at both the fiber and fibril levels and highlighting the potential for broader applications of the multi-scale, multi-modal approach in collagenous tissue studies.

## 1. Introduction

Collagen is the most abundant structural protein in humans and other mammals, and plays an essential role for the structural and functional integrity of many organs [1]. The non-centrosymmetric structure of collagen fibrils contributes to their intrinsic ability to produce strong second harmonic generation (SHG) signals, making SHG microscopy a powerful tool for obtaining high-contrast images of collagen [2,3]. This technique has been extensively used to investigate physiological and pathological conditions that alter collagen microstructure because it can also provide quantifiable measures of anisotropy and disorganization within tissues that are readily interpretable [4–8]. However, another parameter has shown potential to be a diagnostic marker: the forward to backward ratio (F/B), defined as a quantitative measure comparing the intensity of SHG signals emitted in the forward direction through the sample to those reflected backward [9]. The interpretation of the F/B is complex due to its sensitivity to a range of changes in fibrillar structures, including variations in fibril diameter, crosslinking, and organization, and the precise nature of the change causing high F/B is ambiguous [10–20]. Therefore, while SHG excels as a primary method for detecting microscale changes, integrating it with other analytical techniques significantly expands its diagnostic value, providing a more nuanced understanding of specific structural alterations and their implications for tissue health. In this research, we have integrated SHG with nanoindentation (NI) and Atomic Force Microscopy (AFM) to provide a comprehensive analysis of the changes at multiple scales, offering a deeper insight into the structure and mechanics of collagenous tissues including the interpretation of F/B.

Changes in collagen microstructure directly influence the mechanical properties of tissues [21–29]. NI is essential for probing local, microscale mechanical properties, especially for heterogeneous soft tissues [30–33]. The optomechanogram is a co-registration of NI and SHG microstructural organizational parameters, which together provide insights into structure-mechanical property relationship [7]. This method has previously established that higher indentation moduli can result from out-of-plane fibrils in the presence of shear stress induced by indentation [34]. Building on this work, this study explores a novel aspect by introducing the correlation between the F/B from SHG and mechanical stiffness from NI. The significance of this new correlation lies in its potential to reveal that regions exhibiting a higher F/B are also characterized by increased mechanical stiffness. Such findings highlight the potential of this methodological integration as a powerful diagnostic tool for identifying fibrotic conditions potentially induced by environmental exposures or diseases.

While the integration of SHG and NI provides substantial insights into how the mechanical properties of tissues correlate with SHG parameters, it does not clearly demonstrate the underlying cause of the observed variation in F/B. To understand the origins of these changes, we have incorporated Atomic Force Microscopy (AFM) into our study. AFM provides essential nanoscale resolution, enabling the detailed examination of ultrastructural details of fibrillar proteins [35]. SHG excels at visualizing the overall collagen microstructure at the microscale but lacks the resolution for nanoscale insights at the fibril level, crucial for fully understanding the variation in F/B. Although several studies have explored integrating AFM with SHG, with limitations in either not imaging same region across both techniques or the requirement of labelling [36–42]. Our study introduces ‘Targeted Correlative Microscopy’, enabling a combination of AFM and SHG in the same microscopic region, which is essential for characterization of a heterogeneous tissue. This approach not only allows for targeted investigation of variations in the microstructural parameters revealed by SHG but is also straightforward to implement. Such precise integration is essential for pinpointing specific structural alterations at the nanoscale level that may lead to observable changes in SHG or other confocal imaging techniques.

This novel integration of SHG, AFM, and NI establishes a comprehensive framework for studying collagenous tissues. By synchronizing the quantitative analysis of the F/B from SHG with the detailed structural insights from AFM and the mechanical data from NI, we create a multi-scale, multi-modal perspective on collagen architecture within the tissue. This approach allows for a nuanced understanding of the localized changes within tissues, highlighting regions of potential disruption and abnormality that might not be apparent when these modalities are used independently. To illustrate the practical application and efficacy of this methodology, we applied it in a targeted case study examining the effects of an environmental toxicant - propylparaben (PP) on uterine tissue. This application not only showcases the capabilities of our approach but also connects it directly to an important issue in women reproductive health.

Research on the impact of environmental factors on reproductive health is gaining increasing attention. Parabens, which are ubiquitous preservatives found in consumer products such as cosmetics, food, and pharmaceuticals, have been under scrutiny; previous studies have linked paraben exposure to decreased fertility and adverse birth outcomes, including lower birth weight [43–46]. Although there is consensus that parabens act as endocrine disruptors, the specific mechanisms through which parabens may affect uterine function remain poorly understood, highlighting a significant gap in current research. This case study, while not delving into these underlying biological mechanisms, seeks to address some of this gap by investigating the effects of PP, one of the most prevalently used parabens, on the uterus with the integrative approach described above. We hypothesized that exposure to PP leads to measurable changes in the mechanical properties and collagen microstructure of the uterus, which can be detected and quantified through our integrated use of SHG, AFM, and NI. Our objectives were to obtain quantitative parameters from SHG to examine how the chronic PP exposure changes the collagen microstructure, to evaluate the impact of PP exposure on the stiffness using NI, and to interpret the significance of F/B by assessing whether regions with altered F/B exhibit distinct mechanical responses and nanoscale structures, as indicated by the correlation of SHG with NI and AFM respectively.

We observed that PP exposure resulted in a heterogeneous distribution of fibrotic regions within the uterus. These affected regions exhibited higher F/Bs in quantified SHG images, increased stiffness in NI, and alterations at the fibrillar level, including increased fibril diameter and highly entangled collagen fibrils as revealed by AFM. These findings hint at the potential role of PP in affecting reproductive health. This case study not only highlights the specific effects of parabens on uterine tissue but also demonstrates the general applicability of our integrated methodology to assess the impact of environmental toxicants on tissue structure and function at multiple scales.

## 2. Methods

### 2.1. Study design and sample collection

Female CD-1 mice, an accepted animal model in reproductive toxicology studies, were purchased from Charles River Laboratories (Charles River, USA). Mice were housed at the University of Illinois at Urbana-Champaign, College of Veterinary Medicine Animal Facility. Mice were kept under 12-hour light–dark cycles at 22±1°C and were provided food and water ad libitum. Mice were administered daily doses of PP (CAS#94-13-3, Sigma Aldrich, USA) or vehicle control (corn oil) from ages 3 to 9 months, spanning their primary reproductive period [47]. Mice received PP at doses of 2, 20, or 200 µg/kg/day, termed as low, medium and high dose respectively, determined based on human exposure levels reported in NHANES 2015-2016 and estimated daily intake from different countries, adjusted for interspecies differences with a conversion factor of 12.3 [48–50]. Dosing solutions were prepared monthly and administered orally each morning using a pipet tip, a method as described in [51] and chosen for its minimal stress and relevance to human exposure routes; After 6 months, mice were euthanized during the diestrus stage for tissue collection, with one portion of the uterine horn immediately flash-frozen in liquid nitrogen and then stored at - 80°C until further use. The other portion was fixed in 10% neutral buffered formalin (NBF), dehydrated through a graded series of ethanol concentrations (from 70% to 100%), and then embedded in paraffin. The Institutional Animal Use and Care Committee at the University of Illinois at Urbana-Champaign approved all procedures involving animal care, euthanasia, and tissue collection (IACUC# 20034).

### 2.2. Nanoindentation

Frozen uterine horns were thawed, and transverse sections were cut into 2 mm lengths, embedded in an optimal cutting temperature compound, and cryo-sectioned (-20°C) in preparation for both NI and SHG imaging. Keeping several adjacent 10 µm sections aside, a Piuma nanoindenter (Optics11, Amsterdam, Netherlands) with a probe radius of 99 μm and cantilever stiffness of 0.50 N/m, indented the remaining horn tissue while submerged in PBS. Indents were along the medial-lateral line of the uterus at intervals of 50 μm and along two parallel lines spaced 50 µm apart. The maximum indentation depth was set to 5 μm with a displacement rate of 5 μm/s. We evaluated the indentation modulus E* as a measure of local stiffness using the force-indentation data captured by the device and using a generalized anisotropic contact model as described in [52]. Brightfield images of the entire uterine cross-section were captured using the built-in camera of the nanoindenter. The locations at which the nanoindenter probe contacted the tissue were identified on the brightfield images captured by the camera [7].

### 2.3. Second Harmonic Generation (SHG) imaging

SHG imaging was conducted in both forward and backward directions on 10 µm sections (described above) using a confocal microscope (Zeiss LSM 710) equipped with a MaiTai DeepSee laser, which emits 70-fs pulses centered at 780 nm, following the method as described in [12]. The laser beam focused on the sample using a 20x objective with a 0.8 numerical aperture (NA) in order to obtain an image of the entire sample cross-section, with the aid of a motorized stage and image stitching functionality provided by the Zen Microscopy Software. The backward-directed SHG signal was captured using the same objective, while a 0.55 NA objective captured the forward-directed signal. In a similar manner, a 40X, 1.2 NA water immersion objective acquired volumetric images measuring 212x212x10 µm from both directions along the predetermined line of NI (section 2.5). These images were then cropped into 50x50x10 µm sub-volumetric image stacks that were directly along the line of NI for quantification and correlation. The study used 3 mice/dose group (12 mice) and 5 volumetric image stacks per uterine layer, resulting in 15 data sets for each dose and each layer (5 image stacks/layer/mouse x 3 mice/dose group). These images were quantified (section 2.4) and then correlated with the indentation moduli, E*, obtained from the NI experiments (section 2.5). All the measurements used for correlation analyses came from transverse cross-sections. However, we also imaged longitudinal sections to further analyze the microstructure and help to interpret results from the transverse sections.

### 2.4. SHG quantification parameters

Each image in the 50x50x10 µm stack was down sampled in the X and Y directions to make the pixel size equal in X, Y and Z orientations. Two parameters, φ, the out-of-plane angle, and spherical variance (SV), were computed for each of the subregions using orientation data from FT-SHG (Fourier Transform-Second Harmonic Generation) analysis [53,54]. φ is the average angle of the collagen fibers relative to the transverse plane within the volume, where 0° and 90° refer to the fibers parallel to the transverse and longitudinal planes, respectively. SV indicates the dispersion of the collagen fibers in the 3D space, ranging from 0 to 0.5, where 0 indicates that the collagen fibers have a preferred orientation, suggesting that the fibers are aligned in a specific direction, and 0.5 signifies fibers that are randomly oriented with no predominant alignment direction [8]. The F/B is the ratio of the intensity of each pixel in the forward SHG image to its counterpart in the backward SHG image. The F/B was determined for a 50x50 µm region from the central part of each volumetric stack by averaging these pixel-wise ratios. To minimize the influence of noise, only the pixels with considerable signal above the noise floor were included for F/B analysis, using a threshold [12].

### 2.5. SHG and NI correlation

Brightfield images of thick tissue cross-sectional surfaces were captured with both the nanoindenter’s built-in camera and the LSM 710 in brightfield mode, and these bright-field images from both modalities were co-registered manually [7]. Then, a two-step SHG imaging approach was employed to effectively align and correlate the NI and SHG parameters. Initially, SHG was conducted on the indented surfaces of the thick tissues to map and overlay the lines of indentation onto the SHG microstructure. Subsequently, SHG was performed on the adjacent thin section of 10 µm to capture both forward and backward SHG signals (section 2.3) and quantify the SHG parameters (section 2.4) along the pre-established lines of indentation obtained from the initial SHG, enabling correlation analysis between the SHG parameters of interest and the indentation modulus (E*). E* for each subregion was correlated with SHG parameters, using an average of 4 measurements arranged in a square pattern, each separated by 50 µm, to ensure the accuracy of the correlations (Figure S1). In total, n=15 data points per layer per dose and per dose group were used to correlate the indentation modulus with the corresponding SHG parameters of the region.

### 2.6. Atomic Force Microscopy (AFM) with Targeted Correlative Microscopy

Tissues fixed in paraffin were sectioned transversely at 7 µm thickness using a microtome and collected on gridded cover glass with a matrix of marked 500x500 µm grid lines where rows are marked with letters and columns with numbers (Correlative Microscopy Coverslips, 20x20 grid of 0.5mm squares, Structure Probe, Inc.). Paraffin was removed with xylene then samples were dehydrated through a series of ethanol solutions at concentrations of 50%, 70%, 90%, and 100%, each for 15 minutes [35]. SHG was performed to identify regions of interest on the grid for subsequent AFM analysis. For instance, when the 11*K grid square was identified as a ROI using SHG, subsequently, the brightfield objective on the AFM was used to locate the same 11*K grid, and the AFM tip was positioned accordingly (Figure S2). AFM imaging was performed using a Cypher AFM system (Asylum Research, an Oxford Instruments Company, USA) operating in tapping mode to minimize tip-sample forces and preserve the sample integrity. The probes used were Tap300Al-G Tapping Mode AFM Probes with an Aluminum Reflective Coating and a nominal tip radius of 10 nm. Scans were conducted over a 2x2 µm region, with resolution of 11.4 nm maintained consistently across all samples to ensure comparability. We categorized the mice from the control and 20 µg/kg/day dose into 3 subgroups: control, high F/B PP-exposed, and low F/B PP-exposed. For each subgroup, we performed 8 scans, with 2 mice from both the control and the 20 µg/kg/day dose group.

### 2.7. AFM quantification

The AFM software Gwyddion was used for image processing and analysis. Collagen was identified using the extract profile function in Gwyddion, where the trough-to-trough values along the length of the collagen fibrils were measured to check for the periodic bands of the collagen fibrils, which have periodicity of approximately 61-67 nm [36] (Figure S3). Fibril diameters were measured by fitting a Lorentzian curve to line profiles drawn across individual fibrils; diameters were then determined from the full width at half maximum (FWHM) of these fits [36,51,52] (Figure S4). The data for fibril diameters were gathered from n=40 individual fibrils across the 8 scans for each of the 3 subgroups. Additionally, fibril entanglement was assessed by measuring the orientation angles of fibrils relative to the horizontal axis in each scan. The variability of these angles in each scanned region, expressed as their linear standard deviation, provides a quantitative measure of fibril disorganization, reflecting the level of entanglement between the fibrils in that region. The data for fibril entanglement were taken from n=8 scans for each of the 3 subgroups. Unlike automated methods or Fourier Transform (FT) techniques to quantify the entanglement, which might miss some fibrils or incorrectly identify non-collagen structures due to the need for thresholding, this manual approach ensures a more precise and comprehensive assessment of collagen fibril entanglement by avoiding such errors.

### 2.8. Statistical Analysis

Statistical analyses, including principal component analysis (PCA), were performed in R 2023 software (R Foundation for Statistical Computing, Vienna, Austria). A two-way ANOVA with a post-hoc Tukey analysis were used to compare the effects of treatment/dose groups and uterine layers on two parameters, SV and φ. For the E* and the F/B, which did not satisfy the normality assumption, differences across groups were assessed using the Kruskal-Wallis test, followed by Dunn’s post-hoc analysis for pairwise comparisons. Spearman’s rank correlation test was used to evaluate the relationships between structural and mechanical parameters. One-way ANOVA with a post-hoc Tukey analysis was used to compare the fibril diameters and the fibril orientation variability from AFM. A p-value of less than 0.05 was considered statistically significant for all comparisons.

As outlined in section 3.2 of the results, understanding the interaction effect between φ and dose on the F/B was critical. To address that, we employed a linear regression model:

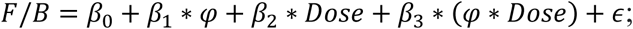

where β0 is the intercept, β_1_ and β_2_ are coefficients for φ and Dose, respectively, β_3_ represents the interaction coefficient between φ and Dose, and ɛ is the error term. The significance of the interaction term was tested through a nested model comparison using ANOVA between two models - one including the interaction term and one without it. This comparison aimed to determine if the addition of the interaction term significantly improved the model fit.

## 3. Results

### 3.1. Backward SHG provides primary insights into layer-specific collagen changes induced by PP exposure

The uterus is composed of three distinct layers: endometrium (Em), circumferential myometrium (CM), and longitudinal myometrium (LM), each characterized by a unique collagen microstructure, as seen in the cross-sectional images of the uterus (Figure 1A). The backward SHG revealed the unique collagen microstructure of the layers in unexposed control uteri, with CM characterized by circumferential in-plane fibrils with low SV (mean ± SEM; 0.10 ± 0.03) and low φ (4.05 ± 2.20); Em characterized by randomly oriented fibrils, with high SV (0.3 ± 0.01) and low φ (8.86 ± 3.38); and LM characterized by out-of-plane fibrils that appear randomly oriented, with high SV (0.33 ± 0.02) and high φ (50.02 ± 3.67).

**Figure 1:**
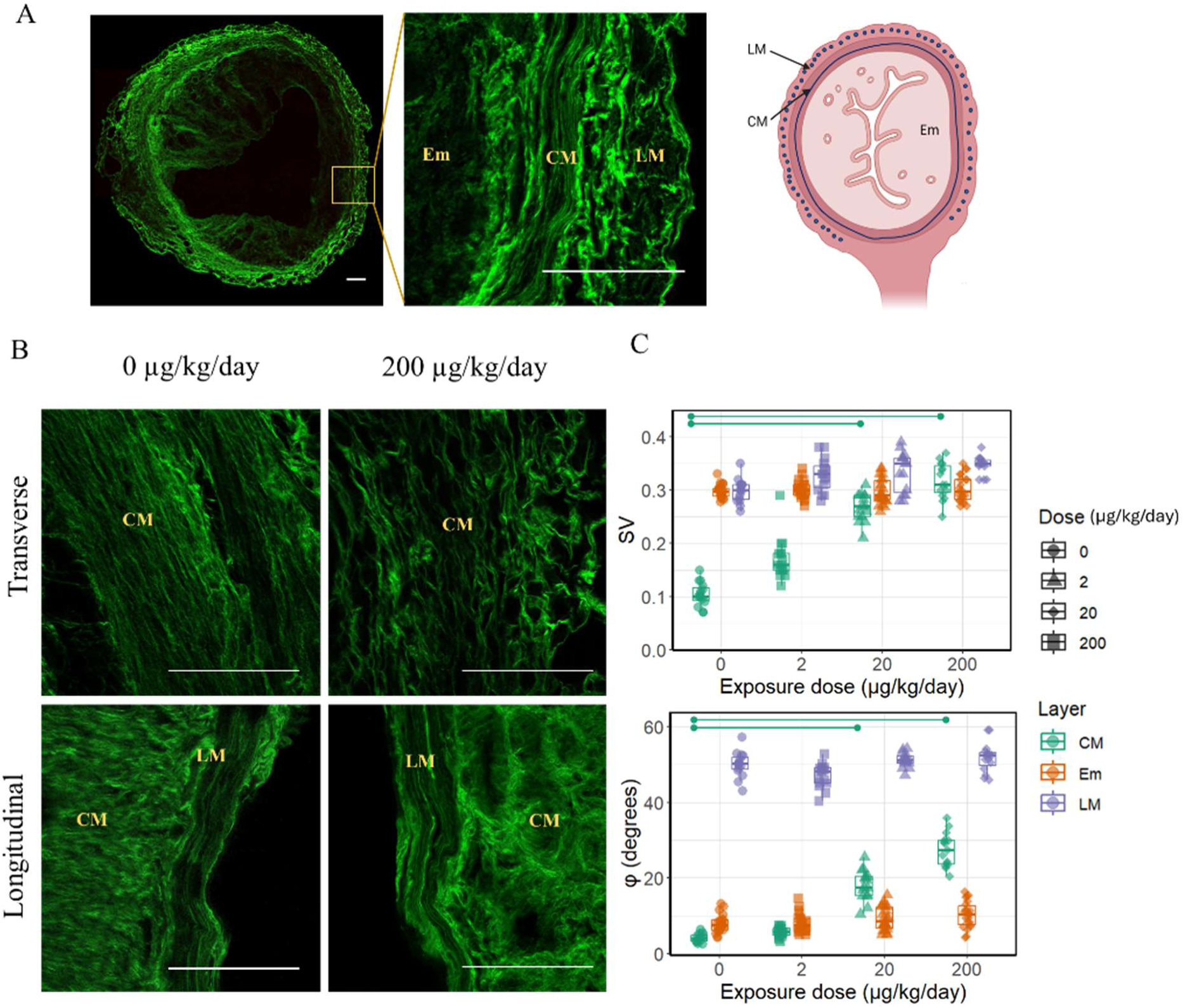
(A) SHG image of uterus transverse section with inset on the right indicating the three layers within the orange box and with schematic transverse cross-section of uterus for reference (Created with BioRender). CM and LM-circumferential and longitudinal myometrium, respectively, Em-endometrium. Scale bar 100 μm. (B) Collagen microstructure in SHG (40X) for control and PP exposed (200 μg/kg/day) uteri in transverse plane (top) and longitudinal plane (bottom). Scale bar 100 μm. (C) SV and φ increase significantly in CM for PP exposure at 20 and 200 µg/kg/day indicating exposure-induced disorganization in the CM layer only (p<0.05).

Backward SHG revealed that the typical circumferential arrangement of collagen in the CM appeared to not be well-preserved in exposed uteri compared to controls, as shown in the transverse sections displayed at the top of Figure 1B. Quantitative analysis showed significant increases in SV (medium: 0.27 ± 0.03, high: 0.32 ± 0.04) and φ (medium: 18.01 ± 4.26, high: 27.47 ± 4.82) with PP exposure, indicating greater disorganization in CM (Figure 1C).

In contrast, LM and Em showed no significant changes in collagen organization post-exposure compared to controls. Since the LM fibrils aligned parallel to the laser might prevent their SHG detection, further imaging along the longitudinal plane was conducted to confirm there were no significant changes, as evidenced by the longitudinal sections shown at the bottom of Figure 1B. Measurements of SV and φ in LM and Em remained consistent and showed no significant differences across all exposure groups, indicating stability in its collagen microstructure despite the exposure (Figure 1C).

### 3.2. F/B revealed further effects of PP exposure that were not evident from backward SHG alone

Expanding our analysis with forward SHG, we noticed an increase in the F/B across all three layers upon exposure, evidenced by SHG images in which red indicates a higher F/B (Figure 2A, 2B for CM and LM, Figure S5 for Em). Figure 2A captures low magnification SHG images from control and PP-exposed uteri at the different doses. These images reveal a heterogeneous distribution of F/B; regions with high F/B exhibit predominantly forward (red) signals, while regions with low F/B display predominantly backward (green) signals. This figure visually demonstrates the overall increase in F/B across exposures and how this change is distributed heterogeneously across different regions in all the layers of the uterus. Figure 2B shows higher magnification SHG images, specifically focusing on the CM of control and exposed uteri from each dose. This figure more closely examines the disorganization within the CM, which is accompanied by high F/B in the medium and high doses, providing a detailed view of the microstructural changes at a higher resolution. Statistical analysis (Figure 2C) confirms statistically significant increases in F/B for medium and high doses (control: 0.07 ± 0.02; medium: 0.93 ± 0.23; high: 0.86 ± 0.10) in CM, (control 0.17 ± 0.03; medium: 1.08 ± 0.18; high: 1.17 ± 0.11) in LM, (control: 0.13 ± 0.03; medium: 0.31 ± 0.12; high: 0.32 ± 0.13) in Em, with no significant changes at the lowest dose for any of the three layers.

**Figure 2:**
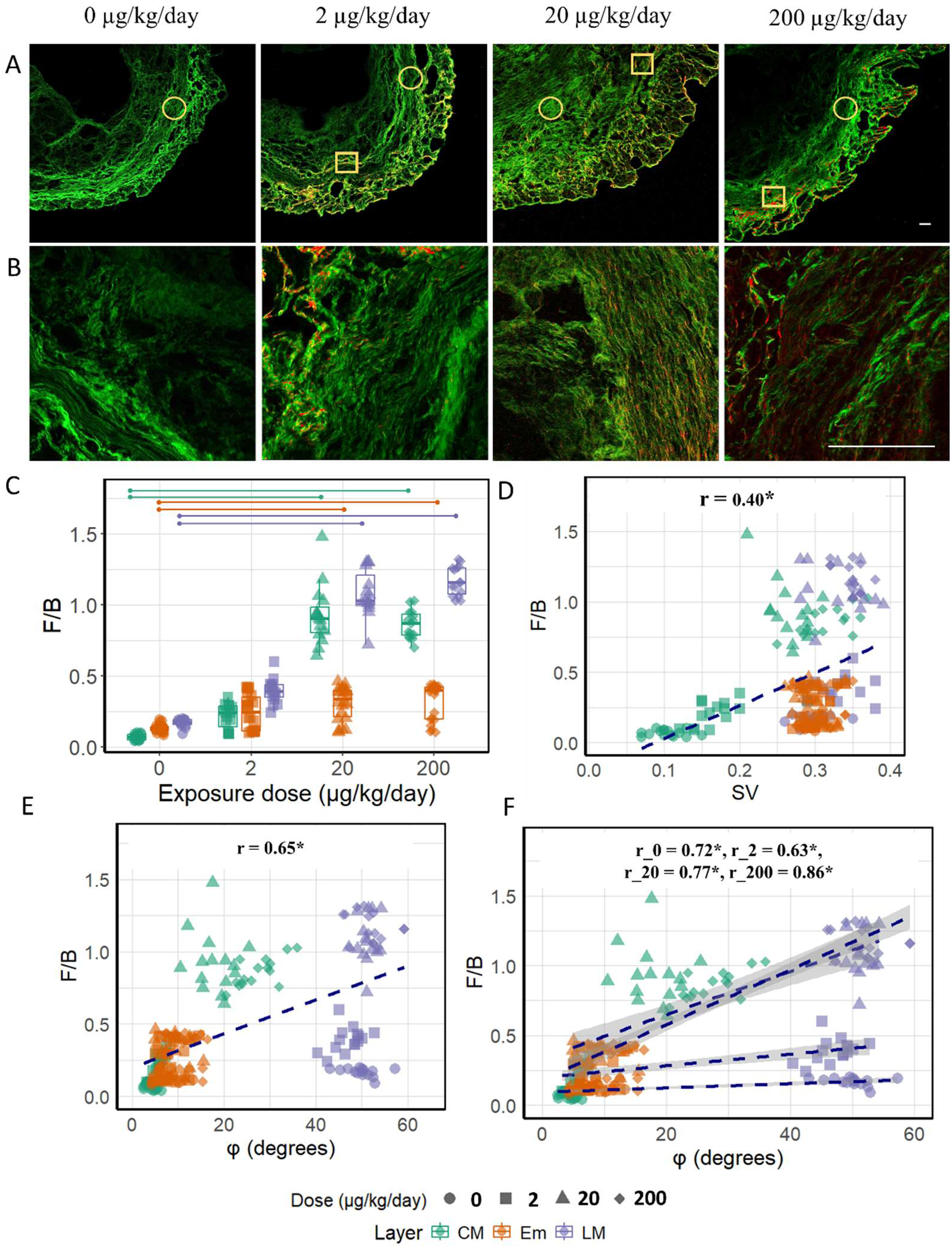
(A) SHG images of control (0) and PP-exposed uteri (2, 20, 200 µg/kg/day) show a heterogeneous F/B distribution in the exposed uteri: high F/B by dominant forward (red) signal and low F/B by backward (green) signals. Representative regions are highlighted: circled regions denote low F/B, while squared regions denote high F/B. Scale bar 100 µm. (B) SHG images of control and PP-exposed uteri show disorganization in CM accompanied by high F/B. Scale bar 100 μm. (C) F/B increases significantly at 20 and 200 µg/kg/day exposure in all layers (p<0.05). (D) F/B vs SV shows a moderate and significant correlation; wide variability in F/B at high SV. (E) F/B increases with φ with all data combined with a strong and significant correlation. (F) F/B increases with φ for each individual dose with significant correlations. For a particular φ, F/Bs are higher in the exposed groups. Also, the sensitivity of F/B to changes in φ is more pronounced in the medium and high-exposure groups, as indicated by the steeper slopes for these groups.

Upon investigating the high F/B across all layers, we aimed to determine if the elevated F/B correlated with shifts in φ or SV to see if the high F/B indicated similar changes that were already captured by SV or φ, or if it suggested additional microstructural changes. Given the potential association between F/B and fibril disorganization [12,14], exploring this relationship was important to accurately interpret the changes. While a moderate and significant correlation (r = 0.40) exists between SV and F/B as shown in Figure 2D, it is notable that higher values of SV are associated with a wider variability in F/B values. This suggests that while there is a general tendency for F/B to increase with SV, the relationship is not predictive as SV increases. In contrast, φ and F/B demonstrated a stronger and significant relationship (r = 0.65), suggesting a more consistent association between these parameters.

Further analysis conducted within each dose group revealed that the F/B ratio increased linearly with increasing φ across all doses. This relationship was marked by strong and statistically significant correlation coefficients: r = 0.72 for the control group, r = 0.63 for the low dose group, r = 0.77 for the medium dose group, and r = 0.86 for the high dose group, as illustrated in Figure 2F. Although this pattern indicates that regions with out-of-plane fibrils exhibit higher F/B in SHG across each dose group, it was also apparent from Figure 2F that for a particular φ, the exposed groups exhibit elevated F/B compared to the control group. This prompted to explore the interaction between φ and dose to understand how they jointly influence F/B.

The nested model comparison using ANOVA, evaluating models with and without the interaction term between φ and dose, revealed a significantly better fit for the model incorporating the interaction term, as reflected by a significant F-statistic of 44.74. This statistic measures how much the model’s explanatory power improves with the interaction term included. This is reflected in Figure 2F, where the exposed groups show an intensified response of F/B to changes in φ compared to the control. For example, a similar shift in φ led to a more pronounced shift in F/B in medium and high PP-exposed groups compared to the control and low PP-exposed groups.

That F/B in exposed groups is notably higher and the responsiveness to changes in φ increases with exposure suggests additional, unobserved changes in fibril structure in SHG imaging that are due to the exposure. These changes could stem from structural modifications within the collagen fibrils, such as thickening or increased crosslinking or nanoscale fibril disorganization, which cannot be observed with SHG and cannot be fully captured by quantitative measurements of φ or SV.

### 3.3. PCA illustrates progressive alterations in uterine microstructure due to PP exposure

PCA was used to further examine the impact of PP-exposure on the microstructure of mouse uterus. PCA is a statistical technique that reduces the dimensions of high-dimensional data by transforming the original variables into a new set of variables. The new variables are the principal components (PCs), which are linear combinations of the original variables and are orthogonal (uncorrelated) to each other. While there can be several, the most important two retain most of the variation present in all the original variables and offer a data representation that reduces the dimensionality [53].

For our study, Figure 3A shows that the first principal component (PC1, Dim1) captures 67% of the variance, while the second principal component (PC2) accounts for an additional 19.6%. The F/B and φ closely align, indicating a correlation. In contrast, SV is nearly orthogonal, suggesting its sensitivity to other aspects of the tissue microstructure.

**Figure 3:**
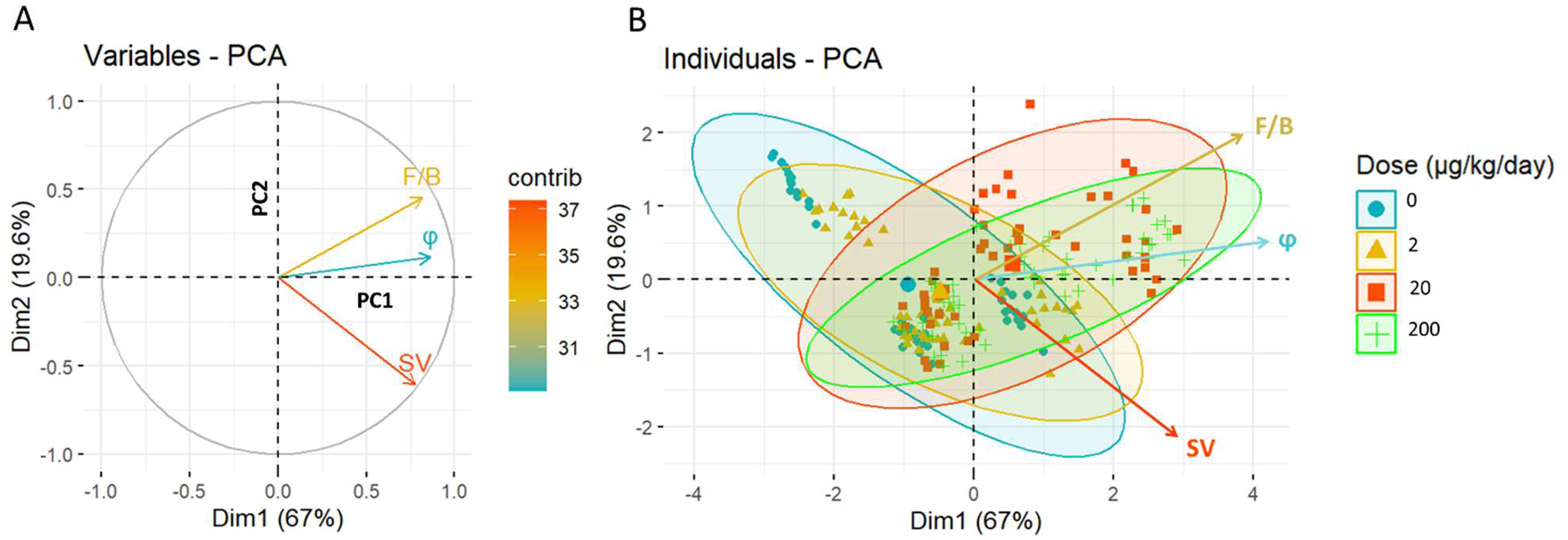
Principal Component Analysis (PCA) of the microstructural parameters. (A) graph of variables showing the relationship between the SHG parameters and the Principal Components (PCs). The angle between vectors shows the degree of correlation; positively correlated variables have adjacent vectors, negatively correlated variables have opposite vectors, and uncorrelated variables have orthogonal vectors. F/B and φ are highly correlated but are independent from SV. (B) graph of individuals in which individuals with similar profiles are grouped together by the shaded ellipses. 95% confidence ellipse is shown for different doses of PP-exposure: control (0), low (2 μg/kg/day), medium (20 μg/kg/day), and high (200 μg/kg/day).

The PCA results (Table 1, Figure 3) demonstrate that F/B, φ, and SV, strongly correlate with PC1, confirming their important contributions to capturing the primary structural changes associated with PP-exposure. While SV aligns closely with PC2 (67%, 0.65)), F/B also shows a moderate yet opposite correlation with PC2 (29%, -0.43), distinguishing itself from φ’s negligible correlation with this component (4%, -0.16). This dual contribution of F/B to both PCs underscores its critical role in capturing a broader spectrum of structural variance within the dataset.

**Table 1.** Correlation between traits and PCs and contributions (%) of traits to the PCs.

| Trait | % Variance Explained by PC1 | Correlation with PC1 | % Variance Explained by PC2 | Correlation with PC2 |
| --- | --- | --- | --- | --- |
| F/B | **33.95** | **0.82** | **29.03** | **-0.43** |
| $\varphi$ | 37.53 | 0.86 | 4.03 | -0.16 |
| SV | 28.52 | 0.75 | 66.94 | 0.65 |

In Figure 3B, the variation in ellipses across different groups indicates significant changes in microstructural parameters upon PP exposure. Notably, the ellipses corresponding to the control and low-exposure groups are narrower along the F/B axis, suggesting a more uniform F/B ratio within these groups. In contrast, the ellipses for the medium and high-exposure groups are noticeably wider along the F/B axis, reflecting a greater variability in F/B. This increased variation in F/B among the higher exposure groups underscores its utility as a critical marker of tissue response to PP exposure. Incorporating F/B in the analysis reveals crucial differences that would otherwise remain undetected, emphasizing the importance of this measure in understanding the impact of PP exposure on tissue microstructure.

### 3.4. NI indicates mechanical consequences in high F/B regions

We observed an increase in E* across all PP-exposed groups, as evident from optomechanograms that highlight the elevated E* in all layers (Figure 4A). The Em is, in general, significantly softer than LM and CM because the constituents in Em are different. Therefore, Em data are shown separately from LM and CM in Figure 4B. In CM, the E* increases from 1.23 ± 0.28 kPa in the control to 6.72 ± 3.52 kPa in the medium and, 7.06 ± 3.73 kPa in the high dose groups. Em increases from 0.2 ± 0.1 kPa in the control to 0.48 ± 0.22 kPa in the medium and 0.54 ± 0.25 kPa in the high dose groups. LM increases from 2.98 ± 0.79 kPa in the control to 11.66 ± 7.13 kPa in the medium and 12.72 ± 4.21 kPa in the high dose groups. The increases are statistically significant only for the medium and high dose groups compared to the control group (Figure 4B).

**Figure 4:**
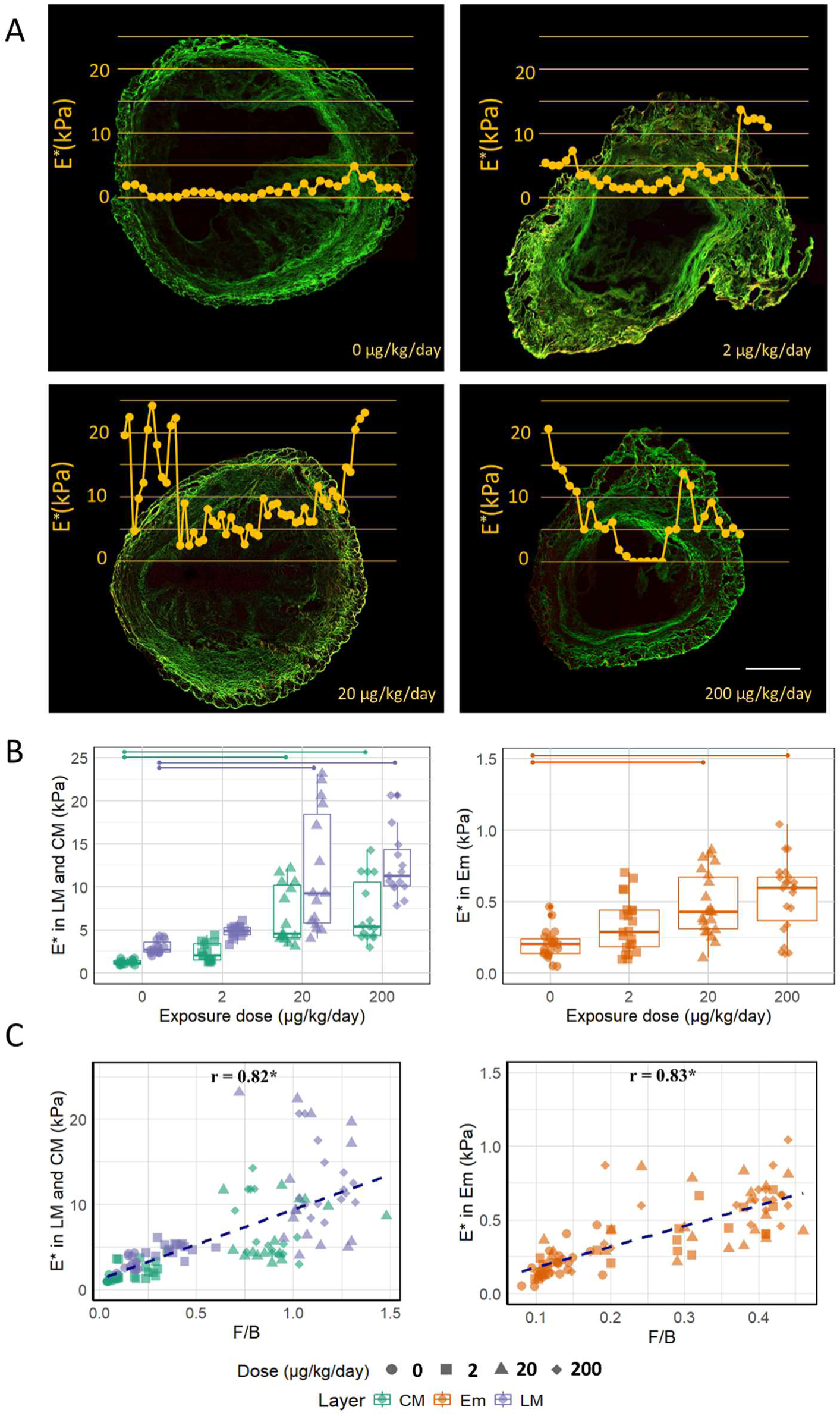
(A) Indentation modulus E* overlaid on SHG images for control (0) and PP exposed uteri (2, 20, 200 µg/kg/day). Yellow lines represent the spatial variation in E* across the uterine layers and they show how E* increases with exposure. The line of indentation coincides with the X axis. Scale bar 500 μm. (B) E* significantly increases from exposure level 0 to 20, 200 µg/kg/day in all layers (p<0.05). (C) E* and F/B have a strong and significant correlation in all the layers.

The longitudinal fibrils in LM show higher E* compared to in-plane fibrils in CM, as expected from our prior research [34]. This increased stiffness is primarily attributed to the fibrils’ capacity to resist shear forces facilitated by the crosslinks along the length of the fibrils. As discussed in section 3.2, PP leads to the re-orientation of in-plane fibrils within the CM, causing them to become more out-of-plane, likely contributing to the observed increase in E* for the medium and high doses. However, despite the lack of significant changes in the out-of-plane orientation of fibrils in the Em and LM, there is still a significant increase in E* in the medium and high doses, suggesting additional changes that stiffen the tissue are at play.

Interestingly, E* demonstrated a high and significant correlation with F/B (0.82 for LM and CM, and 0.83 for Em) suggesting that the factors influencing the increase in F/B in all layers may also be driving the changes in E* (Figure 4C).

### 3.5. AFM imaging reveals nanoscale origins of high F/B regions in the PP-exposed groups

AFM was used to examine collagen fibrils at the nanoscale with the goal of identifying the underlying causes of elevated F/B by observing structural changes not visible with SHG imaging. Three representative 2x2 μm AFM scans of the CM layer of the uterus taken from each of the 3 subgroups-low F/B region from control, and low F/B and high F/B regions from medium PP-exposed group, are shown in Figure 5. Visually, the differences are stark: the control uterus displays thin fibrils with a preferred orientation. Conversely, in the PP-exposed uterus, the low F/B region exhibited disorganized fibrils, like the high F/B region. Yet fibrils in the high F/B region appeared to be notably thicker. This observation prompted a detailed quantification of two key parameters: fibril diameter and fibril entanglement to quantitatively assess the differences observed across the regions.

**Figure 5:**
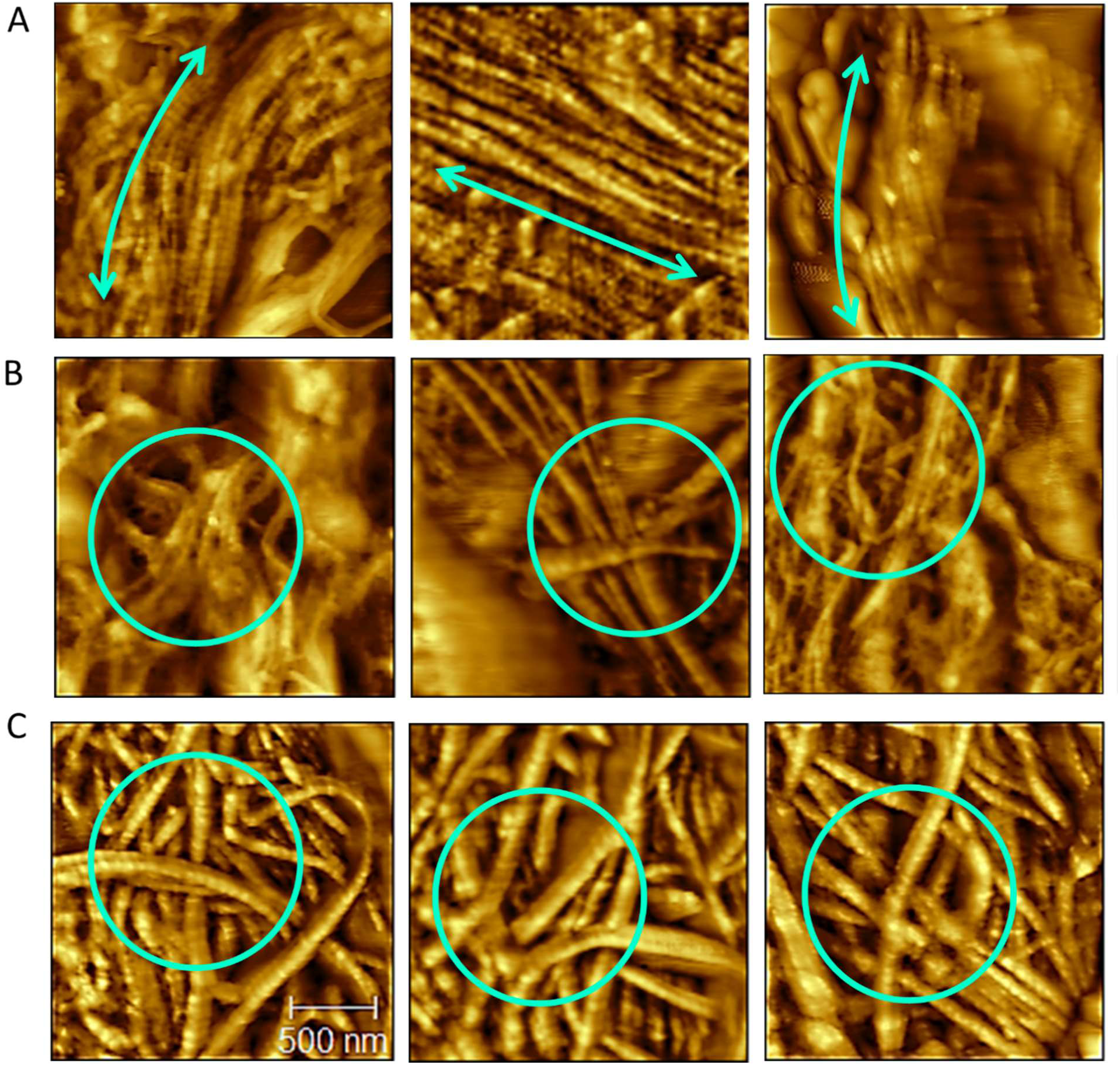
Representative AFM scans of uterine tissue across different conditions. Each panel features three 2x2 µm scans corresponding to each of three groups: low F/B region from control (0 µg/kg/day), low F/B, and high F/B regions from medium exposed group (20 µg/kg/day). (A) control tissues display uniformly thin fibrils, arranged in a preferred orientation (indicated by an arrow), demonstrating an organized structural pattern. (B) Low F/B regions exhibit significant disorganization with irregularly arranged fibrils (highlighted with circle). (C) The high F/B regions, while similarly disorganized, are characterized by noticeably thicker fibrils (highlighted with circles).

Figure 6A illustrates the variations in fibril diameter across the three sub-groups. The control group displayed thin fibrils, which were consistent in diameter (63.6 ± 11.70 nm), indicating a uniform fibril size. The range of fibril diameter in low F/B regions of the PP-exposed tissue (58.9 ± 16 nm) was similar to control. In contrast, high F/B regions of the PP-exposed tissues exhibited significantly thicker fibrils (110.67 ± 25.10 nm) compared to control and the low F/B of PP-exposed tissue.

**Figure 6:**
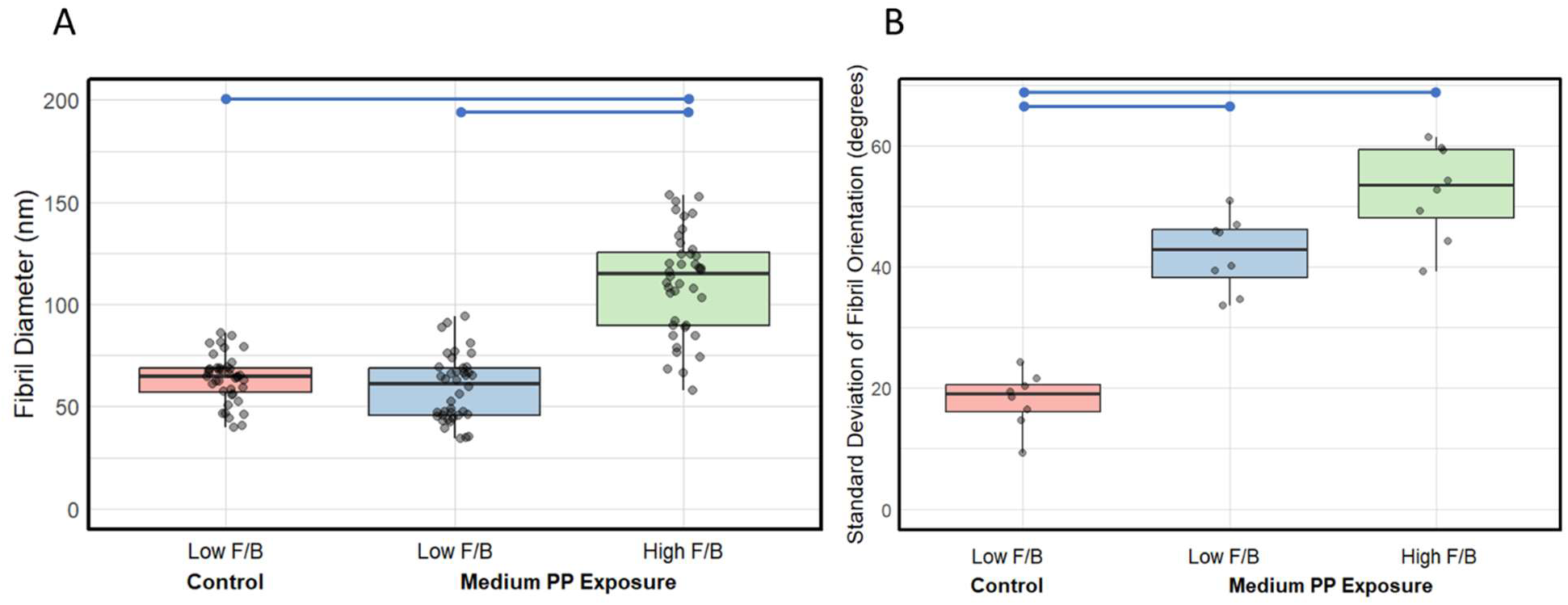
Boxplots showing comparison of the fibril diameter and fibril entanglement of CM layer for the three subgroups: low F/B region from control (0 µg/kg/day), low F/B, and high F/B regions from medium PP exposed group (20 µg/kg/day). (A) Fibrils in the high F/B region have significantly larger diameter compared to those in the control and the low F/B region of the exposed group, with n=40 fibrils for each subgroup. (B) The fibril network in the 2 exposed subgroups displayed significantly higher fibril orientation variability or entanglement than that in the control, n=8 for each subgroup, where each data point represents the variability observed in a single scan.

Figure 6B provides insights into the standard deviation of fibril orientations, a measure of fibril entanglement, across the three sub-groups. Tissue from unexposed mice demonstrated low variability in fibril orientation, consistent with the circumferential pattern of the CM layer (18.1 ± 4.61). In contrast, PP-exposed tissues showed a significant increase in orientation variability, both in low F/B regions (42.2 ± 6.18) and high F/B regions (52.5 ± 7.87), indicative of a substantial increase in fibril entanglement.

## 4. Discussion

The study of collagenous tissue microstructure, especially in response to environmental factors or disease conditions, demands a comprehensive methodology to unravel the complexities at different scales. This work underscores the complementary roles of SHG microscopy, nanoindentation, and AFM in providing a holistic understanding of tissue microstructure and mechanics. Employing this multi-modal and multi-scale approach allowed for an in-depth examination of the structural and mechanical properties of mouse uteri chronically exposed to paraben, serving as a case study to demonstrate the utility and effectiveness of our methods. This integrative approach is novel in its ability to provide a multi-scale perspective that correlates mechanical data with structural imaging on the same tissue regions at different scales, enhancing our understanding of the effect of PP as a model chemical. Notably, our study uncovered significant alterations in the collagen architecture of PP-exposed uterine tissue, marking the first documentation of such changes.

SHG imaging is a pivotal technique for initial assessments of collagen microstructure, particularly useful for quantifying the degree of anisotropy, SV, and the orientation of out-of-plane fibrils, φ. Variations in these metrics, influenced by disease or environmental factors, can signal crucial alterations that potentially impact tissue function. However, there are inherent limitations to consider while interpreting these parameters. Notably, while the uterine layer LM predominantly contains unidirectional longitudinal fibrils, which are inherently anisotropic, when these fibrils are observed from the transverse plane, even slight deviations from their longitudinal orientation can create an appearance of randomness, thus elevating the SV measurements from this perspective. This apparent randomness is a perspective artifact of the transverse section imaging technique and has not influenced the analytical outcomes of this study. F/B is another SHG parameter suitable to assess collagen microstructure, where an elevated F/B potentially indicates undetectable changes at the fibril level. By highlighting regions with higher F/Bs, researchers can efficiently narrow down “regions of interest” for further detailed analysis, making this approach particularly efficient for large tissue samples.

The F/B parameter in SHG, while a potentially powerful diagnostic marker for changes in collagen fibrils, presents several interpretative challenges. This ratio has been used to infer structural changes within tissues, but has been interpreted differently in various studies, underscoring the complexity and sometimes contradictory findings associated with this parameter. AFM correlated with SHG provided the opportunity of looking at the nanoscale structure of the tissue at regions that showed high F/B. The stark differences observed in fibril thickness across control and exposed groups in low and high F/B regions substantiate the hypothesis that the elevated F/Bs detected by SHG are a direct consequence of the thick collagen fibrils. This finding is consistent with several studies that link an increased F/B to collagen fibril thickening [11-17, 20]. However, this finding contrasts with a study reported in reference [10], where no correlation between apparent fiber diameter and F/B was observed, but they relied on SHG to determine the diameter which did not represent the nanoscale changes we observed.

In addition to fibril diameter, we identified from SHG that out-of-plane fibrils also contribute to elevated F/B, which may arise because of the interaction between out of plane fibrils with the excitation laser going ‘through’ the sample, resulting in forward SHG. Contrary to some studies that associate high F/B with fibril disorder [12,14], our findings from AFM in the low F/B subgroup of the exposed tissue suggest that disordered, entangled fibrils which had diameter in the 60-80 nm range did not show high F/B in SHG. F/B has also been shown to be due to the total number of fibrils in a bundle or packing density, which we did not quantify [14, 17] as it was difficult to define a specific bundle. High F/B has been linked to high crosslinking or maturation [19], which we did not verify. Also, the focus was placed solely on the CM layer for this investigation because it contains readily identifiable collagen fibrils, and the other two layers were not investigated. While the F/B serves as a useful comparative tool for assessing tissue changes, its absolute magnitudes can be misleading. The signal intensity is influenced by several factors, including instrument settings and sample preparation, highlighting the necessity of using relative comparisons rather than absolute values.

In addition to underscoring the origin of elevated F/B, AFM played a crucial role in revealing disruptions at the fibril scale. It detected significant fibril entanglement in the PP exposed tissues compared to the organized circumferential arrangement of fibrils in the control uteri. This observation aligns with findings from other researches, which have shown how fibril order can be disrupted in response to any disease or genetic disorders, as detected by AFM in collagenous tissues [57,58].

Nanoindentation results not only showed that PP-exposure increased the stiffness of uterus, but when combined with SHG, it also provided the insight that high F/B regions present in PP exposed tissue are stiffer. This suggests that the fibril thickening observed with AFM contributes to the elevated mechanical stiffness of the high F/B regions, indicating fibrosis. This correlation is pivotal, as it hints that even in the absence of direct mechanical testing tools, the F/B may provide insights into the mechanical integrity of specific tissue regions. It is important to consider that the presence of other components, such as muscle in LM and CM, also influences the observed stiffness levels, and those elements were not investigated in this study. While comparing the two brightfield images from NI and SHG, there is co-registration error involved which should be kept in mind [7]. Also, as it is important for the section to be thin for doing forward SHG [59], the indented surface of the tissue could not be imaged, and the SHG quantifications were done on an adjacent thin section.

The combined use of SHG, NI, and AFM offers a comprehensive set of tools for analyzing collagenous tissues. Each method complements the others, with SHG identifying potential regions of interest across the whole tissue, NI assessing the mechanical implications of microstructural changes, and AFM providing the opportunity to probe the nanoscale structural alterations. However, these techniques do require significant technical expertise and are associated with higher costs, which may limit their widespread application. Moreover, the precise sample preparation and handling needed for these methods can introduce variability in results. Despite these challenges, in the context of imaging techniques, our methodology complements traditional approaches such as MRI, which, while invaluable for providing essential macroscopic and anatomical insights— such as monitoring the structural changes of the uterus during pregnancy [60]—does not capture the detailed micro-and nanoscale structures that can be altered by an environmental or physiological factor. Therefore, our integrated approach not only fills a crucial gap by offering detailed insights at finer scales but also enhances the efficiency of tissue studies by focusing efforts on critical regions. This strategy enriches our understanding of how subtle changes at the microscale can significantly influence broader tissue function and health.

In future studies, we aim to explore how fibrotic changes observed in this study might influence uterine contractility and elasticity, both critical to successful pregnancy outcomes. Further research is needed to correlate these structural changes with fertility outcomes and broader reproductive health. Additionally, we plan to investigate the molecular and biochemical changes accompanying these structural alterations, which may provide deeper insights into the specific reprotoxic effects of paraben exposure.

## Declaration of Competing Interest

The authors declare no competing interests.

## Acknowledgements

This work was supported by NIH1R03ES032887A. Additionally, the lead author was supported partially by Mitul Patel Fellowship. The research was carried out in the Core Facilities at the Carl R. Woese Institute for Genomic Biology, and in the Materials Research Laboratory Central Research Facilities, University of Illinois. We would like to thank Dr. Umnia Doha for suggesting correlative microscope cover glasses. Dr. Amy Wagoner Johnson, Dr. Indrani C. Bagchi, and Dr. Ayelet Ziv-Gal are CZBiohub Investigators.

## Notes

### Competing Interest Statement

The authors have declared no competing interest.

