## Supplemental Material for "Impact of Paraben on Uterine Collagen: An Integrated and Targeted Correlative Approach Using Second Harmonic Generation Microscopy, Nanoindentation, and Atomic Force Microscopy"

**Supplementary figures:**

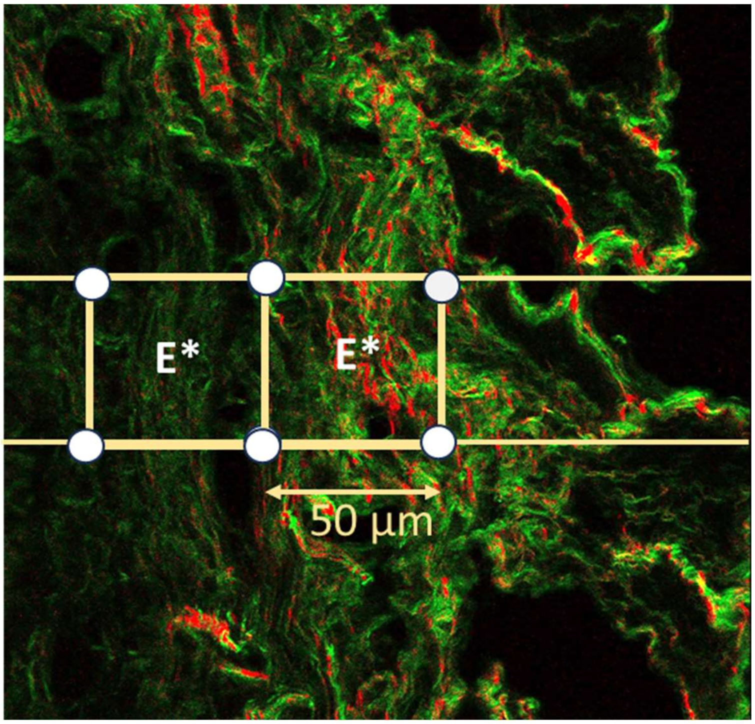


*S1: Methodology for correlating indentation modulus (E*) with the SHG parameters: spherical variance (SV), out-of-plane angle (Phi), and forward/backward (F/B) ratio. E* values were averaged from measurements at the four corners of the displayed 50 µm square. These average E* values were then correlated with the measured SHG parameters of the same square region.*

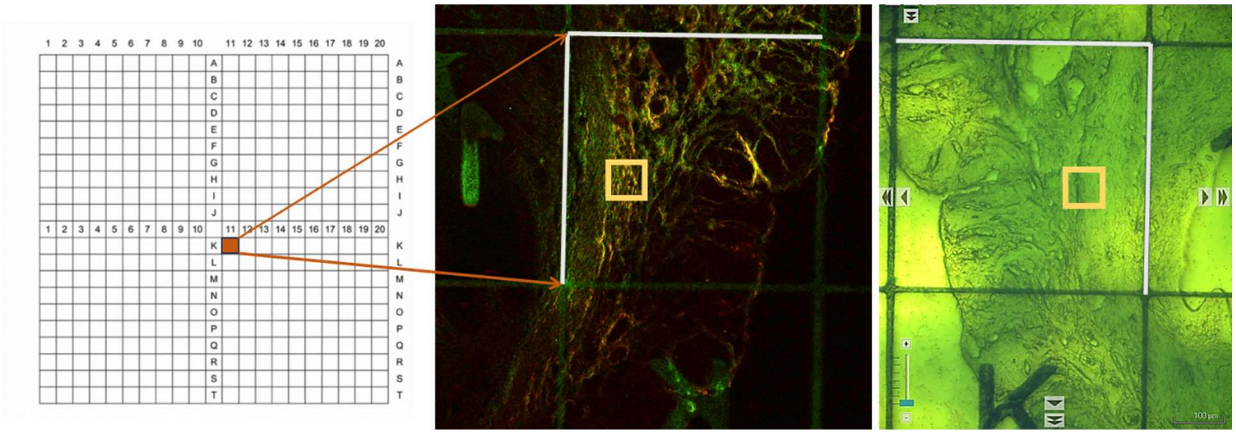


*S2: ROI was first located by selecting a particular grid on SHG, and then the same grid was located using the brightfield objective of AFM. The gridded cover glass was flipped, and the sample was sandwiched between the gridded plastic cover glass and a glass cover glass, resulting in a mirrored view of the grids in the SHG mode.*

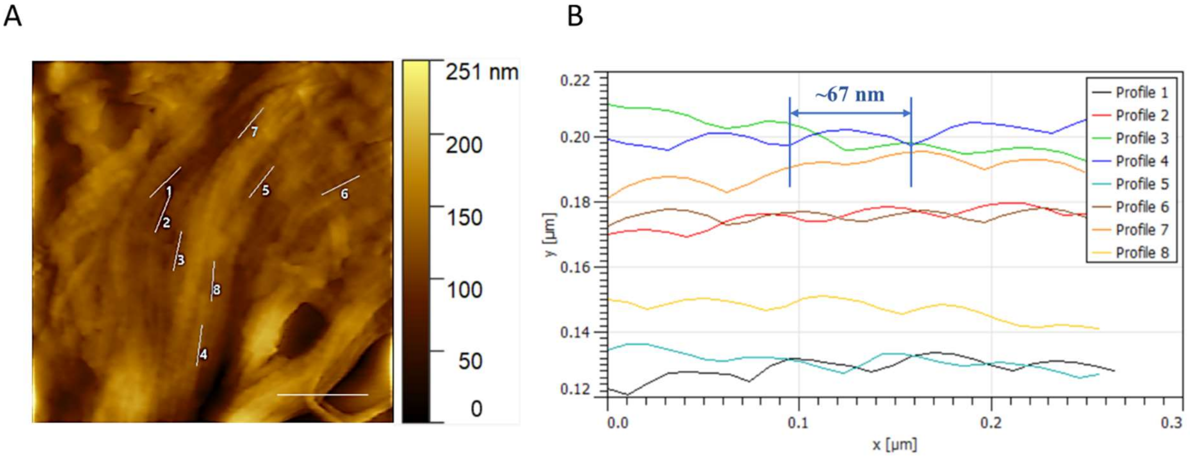


*S3: Quantitative analysis of collagen fibril periodicity. (A) AFM topography of uterine CM layer with 8 representative lines drawn along 8 individual fibrils, scale bar 500 nm. (B) The line profiles along the fibril show a periodicity of ~67 nm.*

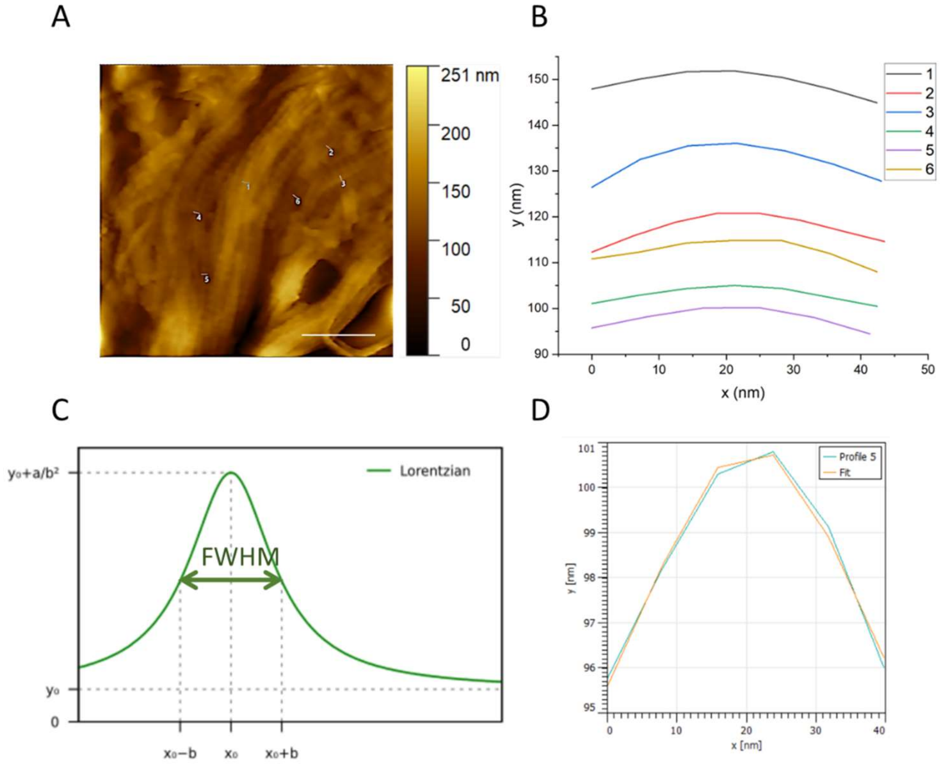


*S4: Quantitative analysis of collagen fibril diameter from AFM. (A) AFM topography of uterine CM layer showing six representative lines drawn across individual fibrils, scale bar 500 nm, with the line profiles on (B). (C) Illustration of the Full Width at Half Maximum (FWHM) on a Lorentzian curve, used to quantify the diameter of a fibril. (D) A representative fit of a line profile fitted with the Lorentzian curve from which FWHM was measured.*

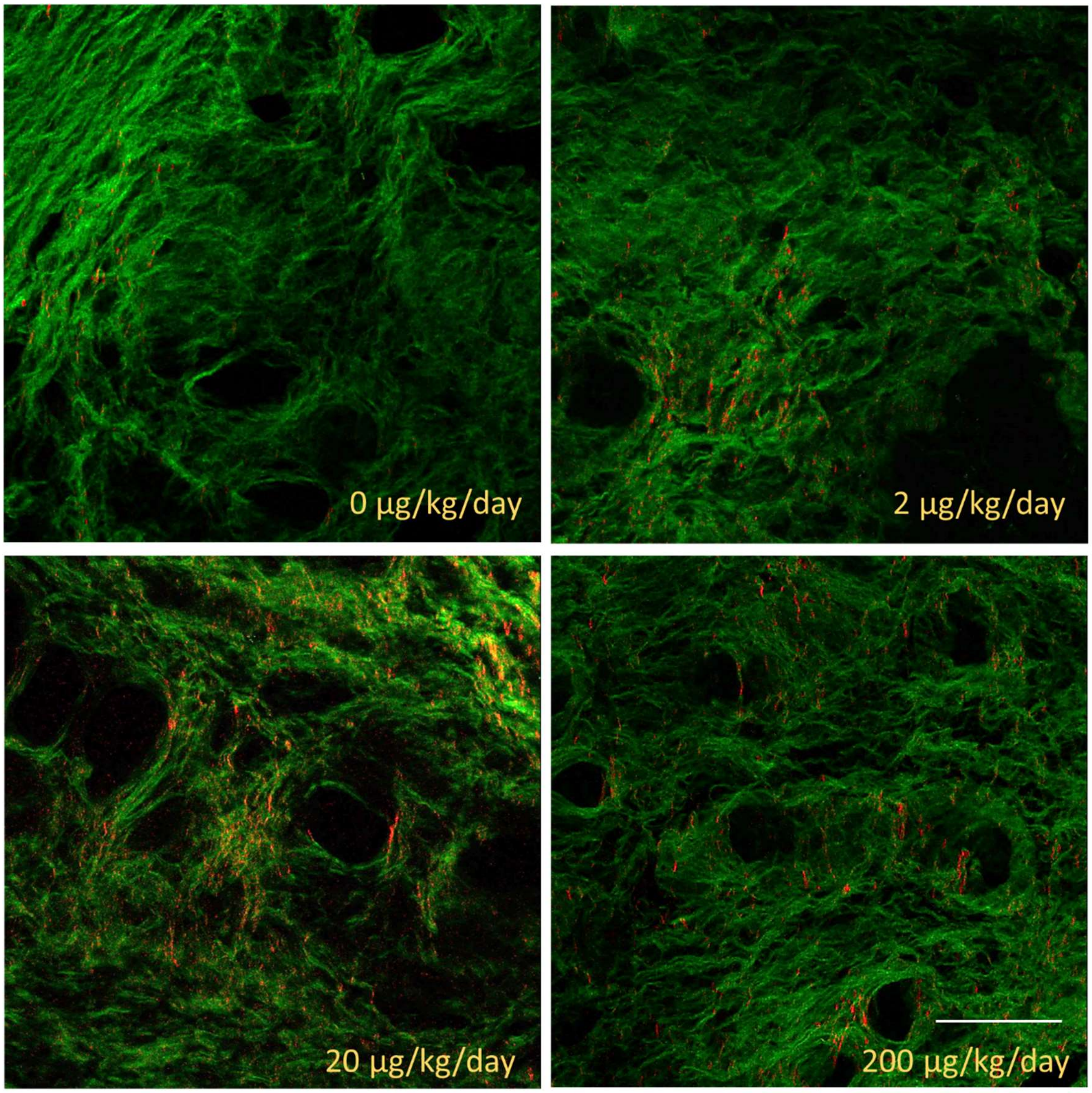


*S5: SHG images of control and PP-exposed uteri show elevated F/B in the Em. Scale bar 50 µm.*
